# CellProfiler: A fit tool for image analysis in droplet microfluidics

**DOI:** 10.1101/811869

**Authors:** Simona Bartkova, Marko Vendelin, Immanuel Sanka, Pille Pata, Ott Scheler

## Abstract

Droplet microfluidic assays are rapidly gaining popularity as the result of the ability to manipulate and monitor single biological molecules, individual cells or small populations of bacteria in pico- and nanoliter droplets, with high sensitivity, precision and accuracy in a high-throughput manner. Nonetheless, there is a demand for user-friendly and low-cost droplet analysis technology. In this article we meet this demand with free open-source software CellProfiler (CP). To illustrate the competence of CP as a droplet analysis tool, we show droplet digital quantification of viable fluorescent bacteria.

## Introduction

Droplet microfluidic assays are increasingly finding the use in microbial and nucleic acid analysis assays. In microbiology, droplet microfluidics has opened up several new experimental possibilities^1^. Examples of microfluidics applications include antibiotic susceptibility studies on population and single cell level^2^, investigation of microbial interactions^3^, and quantification of viable bacteria^4^. Due to its higher precision, sensitivity and robust automatic calibration, droplet digital PCR (ddPCR) has become a strong alternative to traditional quantitative PCR (qPCR) in quantifying nucleic acids in both diagnostics and laboratory research^5,6^. Both, in microbiology and nucleic acid analyses, the detection of positive signals in droplets is usually achieved by measuring their fluorescence^1,7^.

The availability of droplet analysis platforms outside designated microfluidics laboratories is limited. Such limitation commonly reflects the fact that the technology lacks transferability and user-friendliness for non-specialists. Commercial droplet analysis platforms have been developed for ddPCR^5^. Their application can be expanded to include other fluorescent targets like microbes and eukaryotic cells. The platforms provide a combined solution for droplet handling fluidics, fluorescence imaging and data analysis. Bio-Rad combine their QX100/200™ ddPCR™ system with QuantaSoft analysis software^8^. RainDance Technologies® offer the RainDrop *Plus*™ dPCR system and Raindrop Analyst II™ software^9^. In both platforms, fluorescence is read from droplets that are flown in an oil stream one-by-one through the acquisition area in the microfluidic channel^9,8^. Stilla Technologies have a Naica™ System coupled with Crystal Miner analysis software^10^. Here, droplets are imaged together in a monolayer 2D array format^10^. The main disadvantage shared by the commercial systems is their limited accessibility to a wider audience due to high initial hardware and software acquisition costs^5^.

Many microfluidics laboratories have developed their own custom platforms for droplet detection. Similarly to commercial platforms, they are based on either detection of droplet fluorescence in microfluidic channels one-by-one^11,12,13^ or imaging the 2D monolayer of droplets ^14,15^. Both in channel-based and 2D array systems, the droplet fluorescence analysis usually involves developing a custom script based on e.g. Labview ^2,16^, ImageJ^15^, Matlab^17,18^, FluoroCellTrack^19^, or Open Source Computer Vision Library (OpenCV)^20^. However, scripting such analytical tools and customizing them to meet the needs of specific experimental assays in the lab, requires expertise in scripting and programming which is not always sufficiently available in traditional biology and chemistry laboratories.

Here, we argue that there is an unmet need for droplet analysis platforms that are easily adaptable and affordable in conventional chemistry and biology laboratories. For example, depositing droplets as a 2D array on modified microscope slides for imaging is a cost-effective and easy-to-learn approach to introduce high-throughput droplet-based screening for diverse research environments^17^. Suitable programs for the analysis of such arrays are ImageJ^15^ and Matlab^17^, though both of them have a steep learning curve.

In this paper we address this issue and describe a droplet analysis technology based on free open-source software CellProfiler (CP) and its companion CellProfiler Analyst (CPA). The software was created by the Broad Institute Imaging Platform. They developed CP to enable biologists without training in scripting or programming to quantitatively measure phenotypes from high-throughput fluorescence microscopy images^21^. CPA was designed to provide user-friendly tools for interactive exploration and analysis of the data created by CP^22^. This includes a machine learning tool called “Classifier” that can be trained to recognize phenotypes and automatically score millions of cells^22^.

## Results and Discussion

We demonstrate the applicability of our technology using droplet digital quantification of viable fluorescent bacteria. We generate water-in-oil droplets using a microfluidic chip with flow-focusing geometry. Droplets contain diluted bacteria cells, growth media, and the fluorescent tracer dye fluorescein isothiocyanate (FITC) (Fig. 1A). For visualization, we deposit a droplet monolayer on a modified microscope slide and image them with a fluorescence widefield microscope (Fig. 1B).

**Figure 1:**
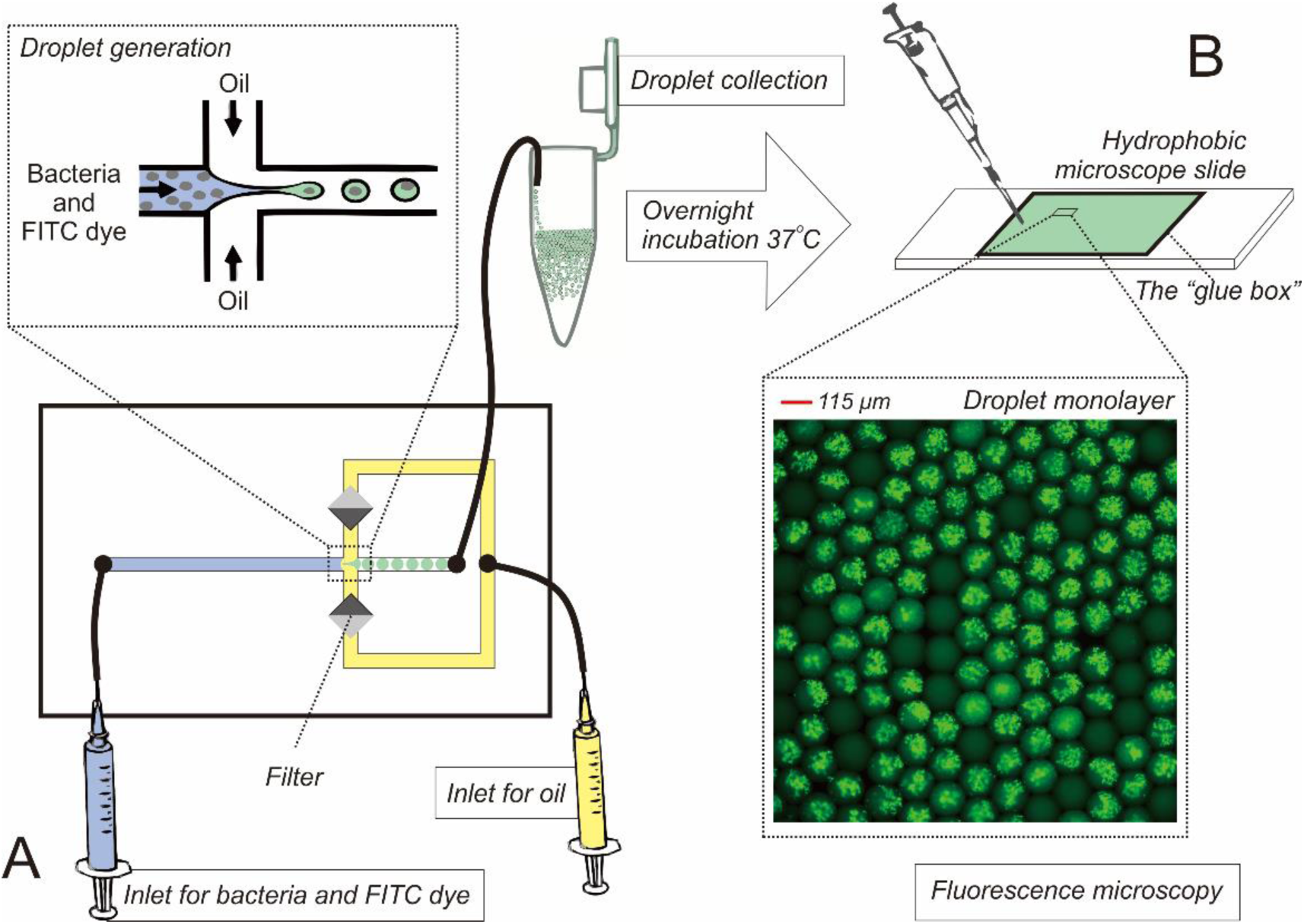
Droplet generation and microscopy. (A) We generate water-in-oil droplets with an average diameter of ∼115 µm, containing bacterial cells, growth media and FITC tracer dye, followed by overnight incubation of droplets at 37°C; (B) Next, we pipette the overnight-incubated droplets on a modified hydrophobic microscope slide where they form a monolayer. Then we fence the droplets by a previously deposited rectangular elevation made of super glue, which we call a “glue box”. Next, we loosely cover the “glue box” with a coverslip. Finally, we visualize the 2D array droplets using a fluorescence widefield microscope. In this experiment, we analyze ∼6900 droplets on a single microscope slide.

We introduce droplet images to our CP image analysis pipeline, by importing raw Tagged Image File (TIF) format images with droplets into the constructed CP pipeline. CP employs Bio-Formats to read input images and can currently read more than 100 available file formats such as BMP, GIF, JPG, PNG, TIF^23^. Lossless formats, such as TIFF, are preferred^23^ and pixels are kept stable, suitable for scientific image data^24^.

The CP pipeline we present identifies droplets and measures their relative fluorescence. We create a pipeline to identify and measure the relative fluorescence intensity of droplets in our images (Fig. 2A). To identify droplets, we use the module “IdentifyPrimaryObjects” (Fig. 2B). This module has several features, including thresholding and object classification, which is usually used for the identification of objects in grayscale images. Conditions for these features are specified by the user. Thresholding turns image pixels into values, which are then classified into two types (black and white). A specific value is set as the limit for each pixel type. This limit is applied for segmentation of objects^25^. CP contains different thresholding methods. In our case, we apply a manual threshold method. This classifies pixels in the background as white color and pixels in the foreground as a black color^26^. We use the module “Images” in CP to visualize the image pixel intensity distribution (Fig. 2C). For guidance to find the best suitable value for our threshold, we use the dip between the histogram peak depicting image background pixels and the peak depicting image droplet pixels (Fig. 2C). Selection of the manual threshold is dependent on the user’s microscope hardware and used labeling techniques. This is why we recommend using the same excitation light and readout sensitivity in practice to simplify analysis. We then apply the “MeasureObjectIntensity” module to measure the relative fluorescence of droplets. Measurement data is automatically saved as either an .h5 file or a .mat file, based on the user’s preference. Module “ExportToSpreadsheet” (optional) exports all measurements as a cvs file, which can be opened in other programs such as Microsoft Excel, R, or python. Finally, we apply the module “ExportToDatabase” to create a SQLite database with all data measurements for further analysis in CPA.

**Figure 2:**
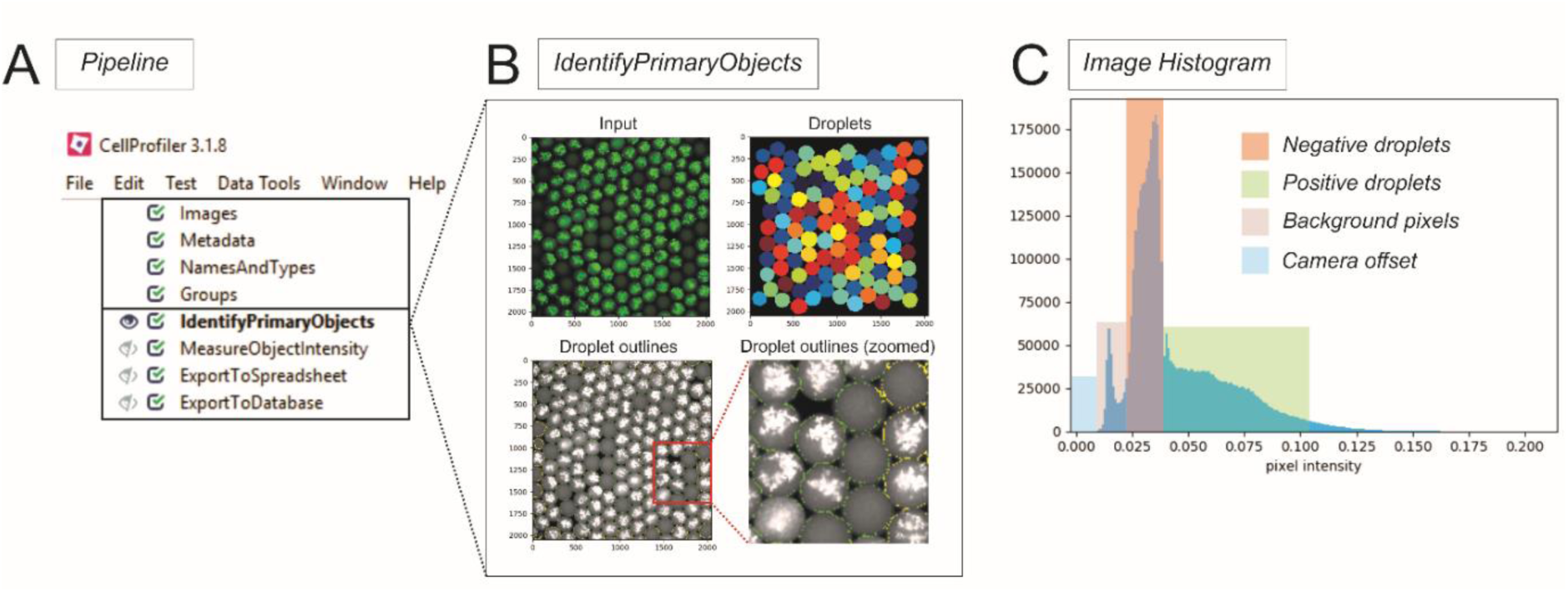
Image analysis with CellProfiler. (A) Overview of our CellProfiler pipeline for analyzing droplet images; (B); We use the CellProfiler module “IdentifyPrimaryObjects” for identifying droplets in each image. The module identifies objects (in our case droplets) based on multiple parameter settings. Key parameters include: (i) typical diameter of objects, (ii) threshold strategy (automated strategies or manual), (iii) method for distinguishing and drawing lines between clumped objects (shape or intensity), (iv) discard objects outside of the diameter range, and (v) discard objects touching the border. We color the “Input image” green for visualization purpose, but all images we use for analysis are greyscale. In the “Droplets outlines” image, all the found droplets with fluorescence intensity units above our threshold have markings around them. Those that are within the correct diameter range are marked with a green outline, while the droplets with a yellow marking are discarded because they are touching the image border; (C) Image histogram from module “Images” depicting four typical pixel intensity sections in our images: camera offset, background, negative droplets and positive droplets.

We use the data generated by the CP pipeline to calculate the number of viable fluorescent bacteria in the sample. We import the data from the CP pipeline stored in SQLite database into CPA, where we i) construct a histogram to display the frequency distribution of the mean of relative fluorescence intensity (Fig. 3A) and ii) employ the CPA Classifier tool to calculate the number of viable fluorescent bacteria in the sample (Fig. 3B). We test accuracy of classification through a confusion matrix, which is a table that describes how well the model for classifying droplets performs based on the training dataset that we construct^22^. Results yield a 99.50% accuracy. We also generate a classification report to evaluate precision, recall metrics, and the F1-score for the two classifications, which results in 1.00 (i.e. best possible score) for both the negative and positive category.

**Figures 3:**
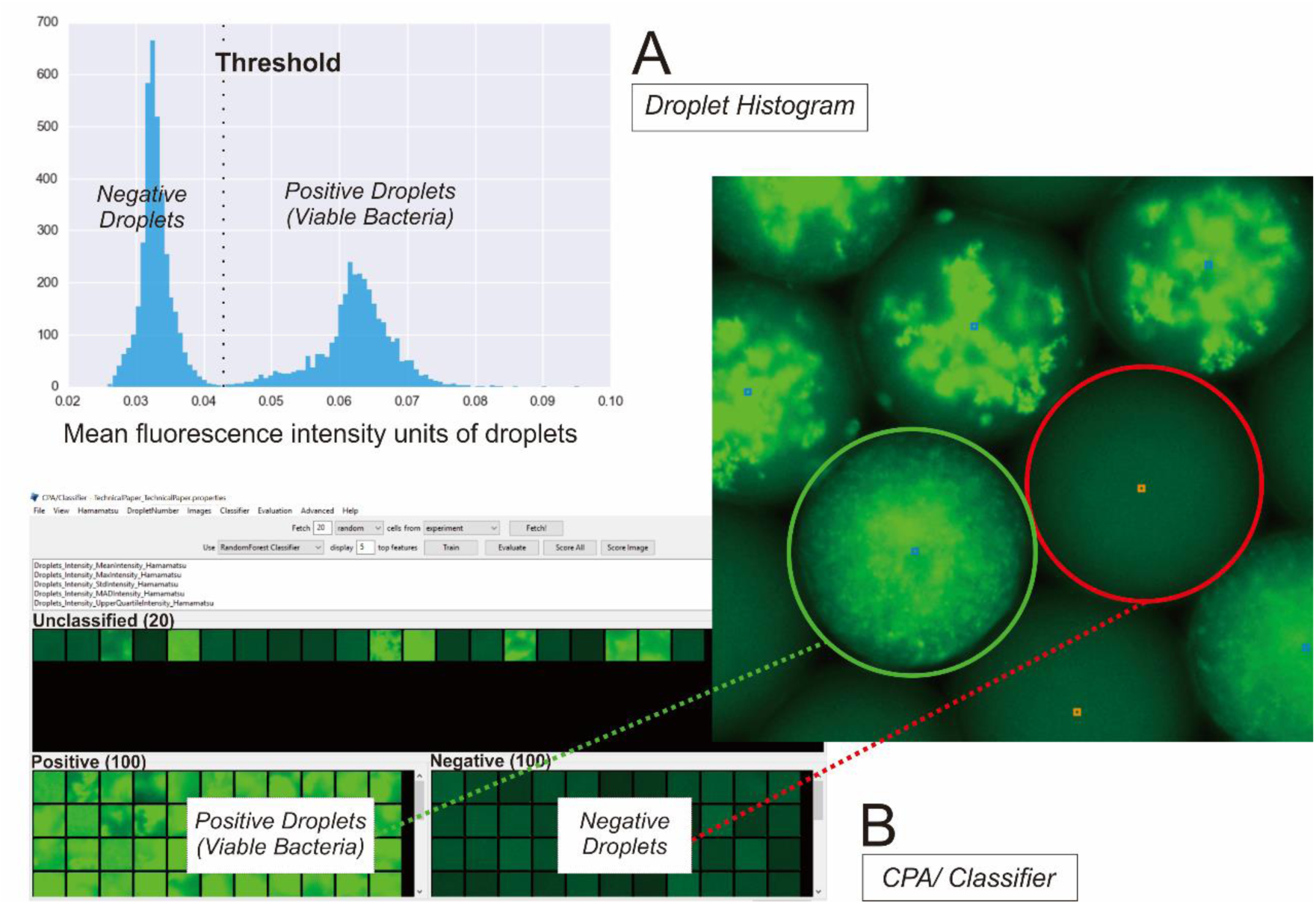
Data visualization and classification with CellProfiler Analyst tools. (A) We visualize data via the “Histogram” tool, based on mean fluorescence intensity units of droplets. For visualization purposes, we draw a manual threshold based on the depicted mean fluorescence between the fluorescence peak of negative and positive droplets, respectively; (B) We train the machine learning tool “Classifier” to automatically classify negative and positive droplets. The overall positive fraction of bacteria is 3153/6899 = 0.45, that translates into mean bacteria cells per droplet λ = 0.61 based on Poisson distribution.

On the basis of our experience, we have found that CP has an intuitive user interface that is easy to learn for general audience. Because of this user-friendly interface, no prior experience in programming is needed for using CP or its companion CPA^27^. Taken together with the available online examples, tutorials, manual and discussion forum, CP is an easy-to-learn analysis software for biologists and other scientists without a background in bioinformatics^27^.

## Conclusion

In this article, we presented a droplet analysis technology based on the free open-source software CellProfiler and successfully analyzed thousands of droplets with viable fluorescent bacteria. The droplet analysis technology we described here is easily adaptable, affordable and automated. This makes it an excellent tool for assisting researchers in conventional laboratories around the world, particularly to extract and analyze quantitative data from digital droplet assays containing thousands of droplets and more. Advantages of this technology include high adaptability, speed, and accessibility at zero cost. The main disadvantage in our case was the inability to import certain file types directly into the software (e.g. Obtained Hierarchical Data Format 5 [HDF5] files). In the end, whether one decides to implement this or another droplet analysis technology, the most critical part is acquiring high-quality images prior to analysis, thereby taking advantage of less manual work, faster analysis and higher adaptability.

## Materials and Methods

### Droplet Generation and Collection

We generate water-in-oil droplets with an average diameter of 115 µm using a poly-(dimethylsiloxane) (PDMS) microfluidic chip with flow-focusing geometry^2^ (Fig. 1A). Droplets contain an overnight culture at 37°C of *Escherichia coli JEK 1036* with chromosome-incorporated gene encoding the green fluorescence protein (GFP), grown in Luria Broth mixed with FITC dye, diluted to roughly 1 bacteria/droplet density and 1µg/mL of FITC (Fig. 1A). We use Novec HFE 7500 fluorocarbon oil with 2% concentration of perfluoropolyether (PFPE)-poly(ethylene glycol) (PEG)-PFPE triblock surfactant for the continuous phase. Surfactant was a kind gift from Professor Piotr Garstecki from Institute of Physical Chemistry, Polish Academy of Sciences. After collection in a 1.5 mL Eppendorf tube, we incubate droplets overnight at 37°C.

### Droplet Imaging

We construct a 1.5 × 1cm rectangular elevation with the height of 250 µm of four layers of super glue (Loctite Precision), which we call the “glue box”, on a microscope slide (Kaltek, ±76 × 26 mm). After drying, we silanize the “glue box” for 3 h under tridecafluoro-1,1,2,2-tetrahydrooctyl-1-trichlorosilane vapors (United Chemical Technologies, USA).

We pipette the droplets onto the hydrophobic Microscope slide where they form a monolayer inside the “glue box”, which we then loosely cover with a coverslip (Kaltek, 18 × 18 mm) (Fig. 1B). Then we acquire fluorescence intensity of bacteria (GFP) and FITC using a fluorescence widefield microscope with the following settings: Objective UPlanFL N 10x/0.3, Excitation LED 470 nm (at 40%), Excitation filter 461–483 nm, Excitation dichronic 488 LPXR, and Emission filter set to 550/88 HC.

### CellProfiler Pipeline

We use CP (version 3.1.8)^21,27^ to design and execute the pipeline. Prior to import into CP, we convert all 64 acquired images in the HDF5 format to grayscale TIF format, due to CP using Bio-Formats to read input images^23^. We import the folder containing the TIF format images into the CP “Images” module. Next we use the “IdentifyPrimaryObject” module to find objects of interest (droplets) in the analyzed images. To find all droplets based on manual examination of 30 randomly selected images from the experiment, we set the threshold manually to 0.023. We enhance image contrast during inspection for visualization purposes, yet use only original images for analysis. Next, we set the typical diameter range of found droplets to 150-250 pixels. All droplets found outside of the diameter range or touching the border of the image are discarded. Separation of clumped droplets is based on the shape. Then we use the “MeasureObjectIntensity” module to measure the pixel intensity units of the objects, with the intensity range set to *Image Metadata* in the “NamesAndTypes”. A detailed explanation of all the available intensity ranges can be found in the CP online manual (http://cellprofiler-manual.s3.amazonaws.com/CellProfiler-3.1.8/index.html). We export the data as a cvs. file to allow optional analysis through other software using the “ExportToSpreadsheet” module. We also export the data to the database SQLite and create a properties file using the “ExportToDatabase” module to enable further analysis of data in CPA. Finally, CP also automatically creates a data output file in the HDF5 format. The pipeline described here is publicly available at our repository on GitHub (https://github.com/taltechmicrofluidics/CP-for-droplet-analysis). An accompanying detailed guide for droplet analysis is available as online supplementary information

### CellProfiler Analyst

We use the properties file created by CP to facilitate access of CPA (version 2.2.1)^22^ to the SQLite database containing the CP pipeline data. We visualize the mean fluorescence intensity units of droplets by use of the CPA histogram tool (Fig. 3A). In CPA machine learning tool “Classifier”, we fetch 200 random droplets from the images and use them to manually create two classification categories: positive (with bacteria) and negative (without bacteria) (Fig. 3B). Each category contains 100 droplets that classifier can use for training. Based on the Random Forest algorithm^28^, classifier acquires the following five top features it utilizes for automatic classification: MeanIntensity, UpperQuartileIntensity, MaxIntensity, StdIntensity, and MADIntensity. We then apply the training set and model to the 64 images to classify all droplets. For future use, we also save the training set as a cvs. file and our model as a .model file. Finally, based on the positive fraction of bacteria, we calculate the mean bacteria cells per droplet using Poisson distribution^2^.

## Supporting information

CellProfiler and CellProfiler Analyst Guide

## Supporting Information

Detailed guide for droplet analysis using our publicly available pipeline (https://github.com/taltechmicrofluidics/CP-for-droplet-analysis) and the software CellProfiler (version 3.1.8) and Cellprofiler Analyst (version 2.2.1) (pdf).

## Author Contributions

The manuscript was written through contributions of all authors. All authors have given approval to the final version of the manuscript.

## Acknowledgements

Most of the work was carried out in the laboratory set up with the support from the TTÜ development program 2016-2022”, project code 2014-2020.4.01.16-0032. We also acknowledge Estonian Research Council grants MOBTP109 (OS) and IUT33-7 (MV).

## References

(1) Kaminski, T. S.; Scheler, O.; Garstecki, P. Droplet Microfluidics for Microbiology: Techniques, Applications and Challenges. Lab Chip 2016, 16 (12), 2168–2187. https://doi.org/10.1039/C6LC00367B.

(2) Scheler, O.; Makuch, K.; Debski, P. R.; Horka, M.; Ruszczak, A.; Pacocha, N.; Sozanski, K.; Smolander, O.-P.; Postek, W.; Garstecki, P. Antibiotic Inhibition of Bacteria Growth in Droplets Reveals Heteroresistance Pattern at the Single Cell Level. bioRxiv 2019, 328393. https://doi.org/10.1101/328393.

(3) Park, J.; Kerner, A.; Burns, M. A.; Lin, X. N. Microdroplet-Enabled Highly Parallel Co-Cultivation of Microbial Communities. PLoS One 2011, 6 (2), e17019. https://doi.org/10.1371/journal.pone.0017019.

(4) Scheler, O.; Pacocha, N.; Debski, P. R.; Ruszczak, A.; Kaminski, T. S.; Garstecki, P. Optimized Droplet Digital CFU Assay (ddCFU) Provides Precise Quantification of Bacteria over a Dynamic Range of 6 Logs and Beyond. Lab Chip 2017, 17 (11), 1980–1987. https://doi.org/10.1039/C7LC00206H.

(5) Cao, L.; Cui, X.; Hu, J.; Li, Z.; Choi, J. R.; Yang, Q.; Lin, M.; Ying Hui, L.; Xu, F. Advances in Digital Polymerase Chain Reaction (dPCR) and Its Emerging Biomedical Applications. Biosens. Bioelectron. 2017, 90, 459–474. https://doi.org/10.1016/J.BIOS.2016.09.082.

(6) Gorgannezhad, L.; Stratton, H.; Nguyen, N.-T. Microfluidic-Based Nucleic Acid Amplification Systems in Microbiology. Micromachines 2019, 10 (6). https://doi.org/10.3390/mi10060408.

(7) Scheler, O.; Postek, W.; Garstecki, P. Recent Developments of Microfluidics as a Tool for Biotechnology and Microbiology. Curr. Opin. Biotechnol. 2019, 55, 60–67. https://doi.org/10.1016/J.COPBIO.2018.08.004.

(8) Hughesman, C. B.; Lu, D. X. J.; Liu, K. Y. P.; Zhu, Y.; Poh, C. F.; Haynes, C. A Robust Protocol for Using Multiplexed Droplet Digital PCR to Quantify Somatic Copy Number Alterations in Clinical Tissue Specimens. PLoS One 2016, 11 (8), e0161274. https://doi.org/10.1371/journal.pone.0161274.

(9) Rutsaert, S.; Bosman, K.; Trypsteen, W.; Nijhuis, M.; Vandekerckhove, L. Digital PCR as a Tool to Measure HIV Persistence. Retrovirology 2018, 15 (1), 16. https://doi.org/10.1186/s12977-018-0399-0.

(10) Madic, J.; Zocevic, A.; Senlis, V.; Fradet, E.; Andre, B.; Muller, S.; Dangla, R.; Droniou, M. E. Three-Color Crystal Digital PCR. Biomol. Detect. Quantif. 2016, 10, 34–46. https://doi.org/10.1016/j.bdq.2016.10.002.

(11) Pekin, D.; Skhiri, Y.; Baret, J.-C.; Le Corre, D.; Mazutis, L.; Ben Salem, C.; Millot, F.; El Harrak, A.; Hutchison, J. B.; Larson, J. W.; et al. Quantitative and Sensitive Detection of Rare Mutations Using Droplet-Based Microfluidics. Lab Chip 2011, 11 (13), 2156. https://doi.org/10.1039/c1lc20128j.

(12) Lim, S. W.; Tran, T. M.; Abate, A. R. PCR-Activated Cell Sorting for Cultivation-Free Enrichment and Sequencing of Rare Microbes. PLoS One 2015, 10 (1), e0113549. https://doi.org/10.1371/journal.pone.0113549.

(13) Berry, S. B.; Lee, J. J.; Berthier, J.; Berthier, E.; Theberge, A. B. Droplet Incubation and Splitting in Open Microfluidic Channels. Anal. Methods 2019, 11 (35), 4528–4536. https://doi.org/10.1039/C9AY00758J.

(14) Bian, X.; Jing, F.; Li, G.; Fan, X.; Jia, C.; Zhou, H.; Jin, Q.; Zhao, J. A Microfluidic Droplet Digital PCR for Simultaneous Detection of Pathogenic *Escherichia Coli* O157 and *Listeria Monocytogenes*. Biosens. Bioelectron. 2015, 74, 770–777. https://doi.org/10.1016/J.BIOS.2015.07.016.

(15) Demaree, B.; Weisgerber, D.; Dolatmoradi, A.; Hatori, M.; Abate, A. R. Direct Quantification of EGFR Variant Allele Frequency in Cell-Free DNA Using a Microfluidic-Free Digital Droplet PCR Assay. Methods Cell Biol. 2018, 148, 119–131. https://doi.org/10.1016/BS.MCB.2018.10.002.

(16) Kang, D.-K.; Gong, X.; Cho, S.; Kim, J.; Edel, J. B.; Chang, S.-I.; Choo, J.; deMello, A. J. 3D Droplet Microfluidic Systems for High-Throughput Biological Experimentation. Anal. Chem. 2015, 87 (21), 10770–10778. https://doi.org/10.1021/acs.analchem.5b02402.

(17) Baccouche, A.; Okumura, S.; Sieskind, R.; Henry, E.; Aubert-Kato, N.; Bredeche, N.; Bartolo, J.-F.; Taly, V.; Rondelez, Y.; Fujii, T.; et al. Massively Parallel and Multiparameter Titration of Biochemical Assays with Droplet Microfluidics. Nat. Protoc. 2017, 12 (9), 1912–1932. https://doi.org/10.1038/nprot.2017.092.

(18) Tamminen, J.; Lahdenperä, E.; Koiranen, T.; Kuronen, T.; Eerola, T.; Lensu, L.; Kälviäinen, H. Determination of Single Droplet Sizes, Velocities and Concentrations with Image Analysis for Reactive Extraction of Copper. Chem. Eng. Sci. 2017, 167, 54–65. https://doi.org/10.1016/j.ces.2017.03.048.

(19) Vaithiyanathan, M.; Safa, N.; Melvin, A. T. FluoroCellTrack: An Algorithm for Automated Analysis of High-Throughput Droplet Microfluidic Data. PLoS One 2019, 14 (5), e0215337. https://doi.org/10.1371/journal.pone.0215337.

(20) Gawryszewski, K.; Rana, Z. A.; Jenkins, K. W.; Ioannou, P.; Okonkwo, D. An Automatic Image Analysis Methodology for the Measurement of Droplet Size Distributions in Liquid–Liquid Dispersion: Round Object Detection. Int. J. Comput. Appl. 2019, 41 (5), 329–342. https://doi.org/10.1080/1206212X.2018.1542555.

(21) Carpenter, A. E.; Jones, T. R.; Lamprecht, M. R.; Clarke, C.; Kang, I.; Friman, O.; Guertin, D. A.; Chang, J.; Lindquist, R. A.; Moffat, J.; et al. CellProfiler: Image Analysis Software for Identifying and Quantifying Cell Phenotypes. Genome Biol. 2006, 7 (10), R100. https://doi.org/10.1186/gb-2006-7-10-r100.

(22) Dao, D.; Fraser, A. N.; Hung, J.; Ljosa, V.; Singh, S.; Carpenter, A. E. CellProfiler Analyst: Interactive Data Exploration, Analysis and Classification of Large Biological Image Sets. Bioinformatics 2016, 32 (20), 3210–3212. https://doi.org/10.1093/bioinformatics/btw390.

(23) Bray, M.-A.; Vokes, M. S.; Carpenter, A. E. Using CellProfiler for Automatic Identification and Measurement of Biological Objects in Images. Curr. Protoc. Mol. Biol. 2015, 109, 14.17.1-13. https://doi.org/10.1002/0471142727.MB1417S109.

(24) Cromey, D. W. Digital Images Are Data: And Should Be Treated as Such. Methods Mol. Biol. 2013, 931, 1–27. https://doi.org/10.1007/978-1-62703-056-4_1.

(25) Singh, K. K.; Singh, A. A Study of Image Segmentation Algorithms for Different Types of Images. Int. J. Comput. Sci. 2010, 7 (5), 414–417.

(26) Singh, R. T.; Roy, S.; Singh, I. O.; Sinam, T.; Singh, M. K. A New Local Adaptive Thresholding Technique in Binarization. Int. J. Comput. Sci. Issues 2012, 8 (6), ISSN (Online): 1694–0814.

(27) McQuin, C.; Goodman, A.; Chernyshev, V.; Kamentsky, L.; Cimini, B. A.; Karhohs, K. W.; Doan, M.; Ding, L.; Rafelski, S. M.; Thirstrup, D.; et al. CellProfiler 3.0: Next-Generation Image Processing for Biology. PLOS Biol. 2018, 16 (7), e2005970. https://doi.org/10.1371/journal.pbio.2005970.

(28) Breiman, L. Random Forests. Mach. Learn. 2001, 45 (1), 5–32. https://doi.org/10.1023/A:1010933404324.

