## Supplementary material for "CellProfiler: A fit tool for image analysis in droplet microfluidics": CellProfiler and CellProfiler Analyst Guide

### CellProfiler and CellProfiler Analyst guide for analyzing droplets

**Start-up**

1. **Download** the software **Cellprofiler (CP)** (version 3.1.8) and **Cellprofiler Analysist (CPA)** (version 2.2.1) here (https://cellprofiler.org/releases/).
2. **Install** the CP and CPA software in your computer.
3. **Open** CP software by double clicking on the CP icon (CPA will be used later).
4. Wait while the program is initialized.
5. A "welcome to CP!" window will appear in the foreground and the CP interface along with the CP terminal for scripting in the background.
6. *Optional*: You can now choose to download and directly load an example pipeline into CP by clicking the Load and example pipeline or click Download a template from the CP website to:
   a. See how the software operates in general with hands-on training
   b. Use an example pipeline for your own image analysis, if they are more suitable for your data.
7. After you are done reading and exploring the "welcome to CP!" window, either click **x** in the corner to remove it or **–** to minimize it.

**CP introduction**

8. Now you can see the interface of CP.
9. When working in CP, it is best to have this window as "full screen".
10. **Pipeline and Modules:**
    a. A **pipeline** is a processing set to be used for your images and consists of various modules that you can add, edit, delete, move etc.
    b. You always start with **4 standard modules** in your pipeline (Images, Metadata, NamesAndTypes, and Groups).
    c. Your pipeline is always displayed to the left.
    d. All modules currently have the √ icon to the left of them indicating that they will be used for the analysis. If you choose to inactivate some of the modules, click the icon by that module and the box will become empty ☐. This means that module will not be active during the analysis.
    e. The module settings for the specific module you are working in is displayed to the right.
    f. When in doubt regarding the meaning of a specific setting, click on the question mark icon **?** located to the right of each setting. Here, they usually give a good explanation of the setting.
    g. When you start adding more modules, they will also have an "eye" icon to the left of the √. This means that when the module has finished analyzing, the result created by this module will be displayed on your screen as a separate window. If you wish not to see the results then click on the "eye" icon and is will be crossed over. *Important note*: During initial testing of a pipeline it is important to always see the results of each module, though when you start your analysis with a large set of images, it is better to have the "eye" icon crossed over at each module and run the pipeline.
    h. You can **add modules** of your choice by *Edit > Add Module* or moving the mouse to the white space where the pipeline is located and right click on the mouse. Or, this can be performed by clicking the + button close to the "adjust module". For deleting the module, the - button can be used after clicking on the module that you want to be removed. Modules can also be moved up and down in the pipeline by dragging with the mouse or by using the ^ and V buttons.

**Pipeline construction/description**

11. **Images module:**
    a. You always start within this module and it is also currently marked with grey color.
    b. Drag and drop your image files and/or folders to the "Drop files and folders here" section.

c. Follow this link https://docs.openmicroscopy.org/bio-formats/5.9.2/supported-formats.html to see what image formats are supported and most suitable for analysis by CP. *Hint:* For image analysis purposes, a lossless format such as TIF or PNG is recommended.
d. *Optional:* You can *Apply filters* for the files you have (e.g. TIF format files only).
e. Double click on an image file to open a separate window and **see the image** and see the pixel intensity values, zoom in, enhance contrast etc.
f. **Zoom in** by clicking the "loop" icon and then making a rectangle with the mouse where you want to zoom in.
g. **Zoom in** even **further** by highlighting another area on the new zoom image.
h. To **move around** in this plane click on the "cross" icon and move around by holding down the right mouse button and moving the mouse.
i. *Important step:* **Measure distances** on the picture (**e.g. diameter of droplets which will be needed later on during for the approximate settings in the FindingPrimaryObejcts module**) by *Tools > Measure length*
    i. Click where you want to start measuring and hold the mouse button down while moving the mouse towards where you want to measure, a red line will appear and the length across that line will appear below in the right corner of the window. Be aware that when you let go of the mouse button, the line and length measurement will disappear.
j. If you go to the image and click with the right mouse button, the following choices will appear:
    i. Open image in new window
    ii. Show image histogram
    iii. Image contrast
    iv. Interpolation
    v. Save subplot
k. **To enhance contrast** click on *Image contrast > Log normalized* and then choose a higher normalization factor than 1.0 (i.e. 1.0 is the current factor for the original image). Be aware that when you enhance the image, it will NOT be automatically saved. You have to click on save, in order to save the enhanced image if you want to keep it.
l. *Critical note:* Do NOT use the enhanced image for analysis in the pipeline.
Example: From the images below, the contrast modification was performed on one image to get a better picture of confined microbes within the droplets and free microbe's droplet. This contrast modification is depicts the actual image, but not is not suitable for image processing material. Notwithstanding, the image may look better (B) than without contrast modification (A), but the pixel intensity is distorted as seen in their respective histograms (C and D). The contrast modification or enhancement may need specific treatment to avoid the distortion, especially for dark image[1]. This would affect the later image processing, including thresholding for segmentation.

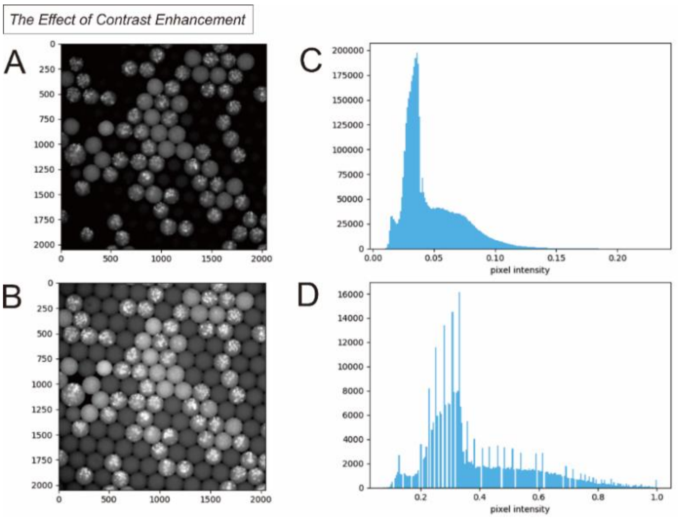

[1] Hussain, Khalid, Shanto Rahman, Md. Mostafijur Rahman, Shah M. Khaled, Abdullah-Al M. Wadud, Muhammad A. Hossain Khan, and Mohammad Shoyaib. 2018. "A Histogram Specification Technique for Dark Image Enhancement Using a Local Transformation Method." IPSJ Transactions on Computer Vision and Applications 10 (1): 3. https://doi.org/10.1186/s41074-018-0040-0

m. *Critical step:* **To see the image histogram** click on *Show image histogram*. Be aware that CP does NOT show you the raw pixel values. The program converts the raw pixel values to values between 0-1 for normalization (see the detailed explanation for this pixel conversion in the CP online manual (http://cellprofiler-manual.s3.amazonaws.com/CellProfiler-3.1.8/index.html). The histogram will look similar to the one depicted and described in our article (Fig. 2C) and the one seen in the above figure (histogram in part C). The **pixel intensity value** corresponding to the slight dip seen between the first small peak and the second big peak is what should be used as an estimate **for the manual threshold** later on in the “IdentifyPrimaryObjects” module. This value will be tested and modified before use on a large image set, nevertheless, it is a good value to start with.

n. **To see the pixel intensity** for a particular spot on the image, simply go to that spot with the mouse pointer without clicking anything and the pixel intensity value of that spot is automatically shown below.

o. You can always click the “house” icon to get the original size image back. However, if you want to see the original contrast of the image back after enhancement you need to go back to the *Image contrast > Log normalized* and write 1.0 as the factor.

12. *Critical step:* **Import** the article **pipeline** by *File > Import > Pipeline from File…*
    a. The pipeline should appear to the left and now include four additional modules:
        i. IdentifyPrimaryObjects
        ii. MeasureObjectIntensity
        iii. ExportToSpreadsheet
        iv. ExportToDatabase
13. *Critical note:* You can add or replace modules in this pipeline if you wish to do so (e.g. if you think you have an uneven background in your images, add modules “CorrectIlluminiationCalculate” and “CorrectIlluminationApply” before “IdentifyPrimaryObjects”). You can find more information about this in the CP online manual (http://cellprofiler-manual.s3.amazonaws.com/CellProfiler-3.1.8/index.html).
14. **Metadata module:**
    a. *Extract metadata?* > “yes”
    b. *Metadata extraction method* > “Extract from file/folder names”. Nevertheless, you can choose another extraction method (e.g. import from file) depending on how you want to extract the metadata from your image files. You can also have several Metadata extraction methods.
    c. *Metadata source* > “File name”
    d. *Regular expression to extract from file name* > “Slide(?P<Site>[0-9][0-9])_”
        i. *Important note:* The exact meaning of each part of the name and how you can edit it to fit your own image file names that CP will recognize, is found by click on the question mark icon **?** to the right.
        ii. We have named our images: Slide01_Bacteria, Slide02_Bacteria, Slide03_Bacteria, etc.
        iii. The expression means that from the image file names (see above section) that you have uploaded in the **Images module**, CP will extract/recognize everything that:
            1. Starts with “Slide”
            2. Continues with capturing the field which we have named “Site”
            3. Continues capturing two numbers between 0-9 (i.e. 2 digits)
            4. Continues again with discarding “_” (i.e. everything that follows is separated from the above)
    e. *Extract metadata from* > “All images”
    f. *Metadata data type* > “text”
15. **NamesAndTypes module:**
    a. *Assign a name to* > “Images matching rules”
    b. *Proceed as 3D* > “No”
    c. *Match* > “All” *of the following rules*
    d. *Select the rules criteria* > “All” *of the following are true*
    e. “File” > “Does” > “Contain” > ”Bacteria”

f. *Critical note:* Text written in CP cannot contain any space between words or the module will not work (e.g. “Fluorescence Image” = Error, “FluorescenceImage” or “Fluorescence_Image” = OK).
g. *Name to assign these images* “FluorescenceImage.” All image file names that contain the writing “Bacteria” thereby be named “FluorescenceImage” in the CP pipeline.
h. *Select the image type* will be set to “Greyscale image”
i. *Select intensity range* will be set according to “Image metadata”
j. You can always click *Update* to see how CP has extracted your images based on your rules to be sure that everything is correct.
k. *Important note:* As mentioned when describing the Images module, CP does not keep the raw pixel intensity values, but converts them to numbers between 0-1. Exactly what it means that the intensity is scaled according to “Image metadata” compared to the other options is explained when you click on question mark icon **?** to the right.
l. *Important note:* If you want to **combine** e.g. fluorescence and bright field **images** or multiple fluorescence images **depicting the same field of view**, you can add multiple names in this module such that they will be analyzed together in the pipeline. If we take our example and say we have both a fluorescence image and a bright field image for each site instead of just one image, we can change the image names to “Slide01_GFP” for the fluorescence image and “Slide01_BF” for the bright field image depicting the field of view we call “01”. Then we change the name in “File” > “Does” > “Contain” > ”**GFP**” and *Name to assign these images* “**GFP**.” We add another name and here we set the name in “File” > “Does” > “Contain” > ”**BF**” and *Name to assign these images* “**BF**.” When you click *Update* is should look like this:

| Update | BF | GFP |
|---|---|---|
| 1 | Slide01_BF.tif | Slide01_GFP.tif |
| 2 | Slide02_BF.tif | Slide02_GFP.tif |

16. **Groups module:**
a. *Do you want group your images?* > “No”
b. We do not use grouping here, since we only have one group of images. However, if you have several groups of images (e.g. experiments on different days.), you can use the grouping and select what you want the grouping to be based upon. More on this can be found in the CP online manual (http://cellprofiler-manual.s3.amazonaws.com/CellProfiler-3.1.8/index.html).

17. **IdentifyPrimaryObjects module:**
a. *Use advanced settings* > “Yes”
b. *Select the input image* > “FluorescenceImage”
c. *Name the primary objects to be identified* “Droplets”
d. *Typical diameter of objects, in pixel units (Min, Max)* > “150 – 250“ pixels for our droplets
e. *Hint:* You can find the approximate diameter in pixels for your own droplets in the Image module. Here you can open an image in another window. By *Tools > Measure length* you can measure the approximate diameter of individual droplets by hand.
f. *Discard objects outside the diameter range?* > “Yes”
g. *Discard objects touching the border of the image?* > “Yes”
h. *Critical step: Thresholding strategy* > “Global”
    i. This strategy is fast and robust and good to use when the background is relatively uniform. If this is not the case, then it is better to use “Adaptive strategy” or use the module “CorrectIlluminiationCalculate” and “CorrectIlluminationApply” before “IdentifyPrimaryObjects.”
i. *Critical step: Thresholding method* > “Manual”
    i. There are several methods for calculating threshold automatically, or you can chose manual and enter a number manually between 0 and 1 for the threshold. Explanation of each method is found when you click on question mark icon **?** to the right.
    ii. Both the automatic and manual options have advantages and disadvantages. For our images, the manual method yielded the best results.

- j. *Critical step:* *Manual threshold* > “0.023” for our images
- k. *Two-class or three-class thresholding* > will disappear as a setting due to the chosen Manual thresholding
- l. *Threshold smoothing scale* > will disappear as a setting due to the chosen Manual thresholding
- m. *Threshold correction factor* > will disappear as a setting due to the chosen Manual thresholding
- n. *Lower and upper bounds on threshold* > will disappear as a setting due to the chosen Manual thresholding
- o. *Method to distinguish clumped objects* > “Shape”
- p. *Method to draw dividing lines between clumped objects* > “Shape”
- q. *Automatically calculate minimum allowed distance between local maxima?* > “Yes”
- r. *Speed up by using lower-resolution image find local maxima* > “Yes”
- s. *Fill holes in identified objects?* > “After declumping only”
- t. *Handling of objects if excessive number of objects identified* > “Continue”

18. **MeasureObjectIntensity module:**
    - a. *Select an image to measure* > “FluorescenceImage” *(from NamesAndTypes)*
    - b. *Select objects to measure* > “Droplets” *(from IdentifyPrimaryObjects #5)*
19. **ExportToSpreadsheet module:**
    - a. *Critical Note:* Export modules always have to be located at the end of a pipeline, because they are based on the data created by the prior modules within the pipeline (i.e. the pipeline always runs step by step, sequentially employing one module at a time.
    - b. *Select the column delimiter* > “Comma (“,”)
    - c. *Output file location* > “Default Output Folder” (or choose any other folder in you wish the files to be exported to).
    - d. *Add prefixes to files names?* > “No”
    - e. *Overwrite existing files without warning?* > “No”
    - f. *Add image metadata columns to your object data file?* > “No”
    - g. *Representation of Nan/Inf* > “NaN”
    - h. *Select the measurements to export* > “Yes”
    - i. *Critical step:* *Press button to select measurements* > Press the button and the following three categories with subitems appear in a new window, where you can choose which measurements to export depending on your experiment:
        1. *Droplets*
        2. *Experiment*
        3. *Image*

        - ii. Each of the three categories will be exported in their own cvs file.
        - iii. The *Experiment* and *Image* categories are always present and give you details regarding the CellProfiler pipeline and the input images respectively.
        - iv. The *Droplets* category is based on the MeasureObjectIntensity module and is the most important category. Here you chose which intensity measurements from each of your droplets in all images you want to export. To be safe choose > “All.”

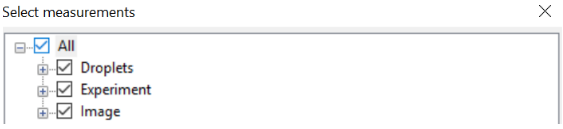

        - v. Also, if more measurement modules would have been added in the pipeline, there would have been more categories.
    - j. *Calculate the per-image mean values for object measurements?* > “No”
    - k. *Calculate the per-image median values for object measurements?* > “No”
    - l. *Calculate the per-image standard deviation values for object measurements?* > “No”
    - m. *Create a GenePattern GCT file?* > “No”
    - n. *Export all measurement types?* > “Yes”
20. **ExportToDatabase module:**
    - a. *Important note:* This module is important if you want to **analyze** all your data **in CPA** after you have run the pipeline. The names you choose to for the files you can chose yourself based on your experiment.

b. Critical step: *Database type* **>** "SQLilte"
    i. This database can handle quite a large dataset and can be stored on a computer or in a larger database of your choice. It is a simpler version of a SQL database that one can use without having to worry about downloading any extra software. Later, MySQL can be used instead if the dataset becomes too big for SQLite to handle).
c. *Experiment name* > "FluorescenceDroplets"
d. *Name the SQLite database file* > "FluorescenceDroplets.db"
e. *Overwrite without warning* > "Never"
f. *Add a prefix to table names* > "Yes"
g. *Table prefix* > "FluorescenceDroplets"
h. Critical step: *Create a CellProfiler Analyst properties file* > "Yes"
    i. This is a text file that tells CPA where to find and extract the data/result from the database that you create)
i. *Which objects should be used for location* > "Droplets"
    i. Tools in CPA can be used to display and classify objects that were identified in the CP pipeline. This setting determines which object centers will be used as the center of the objects that CPA displays. Since we only identified one object (droplets), you only have one choice here. Though, if more objects were identified, then this would be a critical step.
j. *Access Cellprofiler Analyst image via URL?* "No"
k. *Select plate type* > "None"
l. *Select the plate metadata* > "None"
m. *Select the well metadata* **>** "Site"
n. *Include information for all images, using default value?* > "Yes"
o. *Do you want to add group fields?* > "No"
p. *Do you want to add filter fields?* > "No"
    *i.* If you have several experiments or conditions you want to measure, then select "Yes" instead. This will allow you to filter based on your below choices during data analysis in CPA (e.g. filter by date of experiment, thus, only results from that date are shown for a specific analysis).
    *ii.* *Select the plate metadata* > Go back and change this setting from "None" to "Plate"
    iii. When you select yes, another two settings will also appear where you also select:
        1. *Automatically create a filter for each plate?* > "Yes"
            a. The category *plate* is based upon your settings in module" Metadata". In that module you can either add *plate* in the existing extraction method or add another Metadata extraction method for *plate.* Then in module "Groups" you can select "Yes" to grouping and group e.g. based on *plate.*
        2. *Add another filter* > This depends on whether you want to add more categories that you can filter during your data analysis in CPA (e.g. incubation with different substrates or concentrations).
q. *Select the classification type* > "Object"
r. *Enter a phenotype class table name if using the classifier tool in CellProfiler Analyst* > "PosNegDroplets" (or chose another name that you want for classifying your droplets).
s. *Create a CellProfiler Analyst workspace file?* > "No"
t. *Output file location* > "Default output folder" (or choose any other folder)
u. *Calculate the per-image mean values for object measurements?* > "No"
v. *Calculate the per-image median values for object measurements?* > "No"
w. *Calculate the per-image standard deviation values for object measurements?* > "No"
x. *Export objects relationships?* "Yes"
y. Critical step: *Create one table per object, a single object table, single object table or a single object view* > "Single object table"
    i. This way, a single database table will be created that records the object (droplet) measurements. Each row of the table will have measurements for all objects that have the same image and object number.

    ii. You can find more detailed explanation on all choices in the question mark icon **?** to the right.
  z. *Maximum # of characters in a column name* > "64"
  aa. *Write image thumbnails directly to the database?* > "No"

21. **View output Settings:**
    a. Below your pipeline you will find an option to *View output settings*.
    b. Default Input Folder > Choose a folder
    c. Default Output Folder > Choose a folder
        i. This is also where your cvs file from module "ExportToSpreadsheet" and files from module "ExportToDatabase" will be exported to if you chose *Default output folder* as the file location in those modules)
    d. Output Filename > Chose a name
    e. Output file format > HDF5

**Test pipeline in "Test Mode"**

22. **Always test your pipeline on several images**, one by one, before applying the pipeline to the entire image set.
23. Go into test mode by clicking "Start Test Mode"
24. In test mode:
    a. CP goes straight into the first added module in the pipeline, which is also highlighted by CP.
    b. In our case this is the "IdentifyPrimaryObjects" module, where we want to see if our droplets are found correctly.
    c. A yellow warning sign will also appear besides the two Export modules. This just means that these two modules do not work during test mode, since you cannot export anything. Once you exit test mode, the warning goes away.
25. Now you have two choices:
    a. **Click on "Step"** to run the module which is highlighted by CP (i.e. "IdentifyPrimaryObjects") on the first image in your imported image set. This is mode of testing which particularly good when you are interested in testing one of the modules. When you do this:
        i. CP will automatically move on to the next module and highlight it.
26. A new window will also appear with the results from the module (i.e. "IdentifyPrimaryObjects").
27. All the tools and options that were available when you opened an image in "Images" module are also available here (e.g. zoom in and move around to see how well the droplets are identified according to your settings, or enhance the image contrast for better visualization of the droplets).
        i. If you are **satisfied with your results:**
            1. Click on "Step" again and another window will appear with results from the second module (i.e. "MeasureObjectIntensity").
            2. This will show a table with the mean, median and standard deviation of various pixel intensity measurements for your droplets on the image that were found in the "IdentifyPrimaryObjects" module. Here, you will not see the measurements for each individual droplet; however, when you export your results later, you will have measurements for each individual droplet that was identified.
        ii. If you are **NOT satisfied with your results:**
            1. Go back and click on "IdentifyPrimaryObjects" again
            2. Edit the settings in the module and click on "Step" again.
            3. The old window with results will then be replaced by a new one with the latest results.
            4. Repeat this step until you are satisfied with the results.
    b. **Click on "Run"** to run the entire pipeline (i.e. "IdentifyPrimaryObjects" and "MeasureObjectIntensity") on the first image in your imported image set. This way the first image will immediate go through both modules and new windows with results from the modules will appear. This is a faster mode for testing the entire pipeline at once, but, it can become confusing if you have many modules in your pipeline.
28. Now **check more images**. This can be done in several ways:
    a. Click on "Next Image Set"
    b. *Test > Next Image Set*

c. *Test > Choose Image Set*
d. *Test > Random Image Set*

29. Once you are satisfied with your tests and the pipeline, then you can click on "Exist Test Mode".
30. Now you are ready to analyze you entire image set in the pipeline by clicking "**Analyze Images**".

**Data analysis options**

31. Once finished with your analysis, you have several ways to analyze your data:
    a. Analyze the entire data set at once by opening the **cvs file** in **Microsoft Excel, R, Python** etc.
    b. Analyze single images directly in **CP** via "**Data Tools**" by importing the created **HDF5 file**.
    c. Analyze the entire data set at once by opening **CPA** and importing the created **properties file**. *Important note:* Here we will continue with this option and analyze droplets with CPA.

**Data analysis with CPA (version 2.2.1)**

32. **Open** CPA software by double clicking on the CPA icon.
33. **Import** your **properties file**, which you created along with your database in the "ExportToDatabase" module.
34. Once imported, CPA interface and terminal will open, including eight data analysis tools:
    a. Image Gallery
    b. Classifier
    c. Plate Viewer
    d. Scatter Plot
    e. Histogram
    f. Density Plot
    g. Box Plot
    h. Table View
35. In this procedure we will only focus on two of the tools: Histogram and Classifier. If you are interested in other tools or a more detailed explanation of CPA in general, you can find more information in the online manual for CPA (version 2.2.1): https://cellprofiler.org/cpa/

**CPA Histogram**

36. To visualize our droplet pixel intensity measurements made through the CP pipeline, we use the histogram tool.
37. Click on the Histogram icon.
38. An empty histogram plot will appear with several setting options.
39. We want to visualize the mean droplet pixel intensity (though you can visualize any measurement you like).
40. This is done by choosing:
    a. X-axis: FluorescenceDropletsPer_**Object**
    b. X-axis: Droplets_Intesnity_**MeanIntensity**_FluorescenceImage
    c. *Optional:* Change the number of bins you want in your histogram, though 100 as the default is usually ok for the display.
    d. *Optional:* If you want some filters, you can choose one in the filter option.
    e. Then click "Update Chart" to visualize the histogram.
    f. You can save your histogram as an image in various formats.

**CPA Classifier**

41. Here you can train CPA to classify your negative droplets and positive droplets in your entire image set.
42. You can also add new classes and classify droplets in several classes based upon your goals.
43. *FluorescenceImage* > "Green" since it is visually nicer to see fluorescence in green than grey.
44. Click "Fetch!"
45. You now have 20 unclassified objects/droplets randomly chosen from the entire image set.
46. **To enhance the contrast** of the droplet for enabling better classification, go to *View > Image Controls* which gives you a new window where you can enhance the contrast.

47. Each square represents the center of a droplet and if you move to a square with the mouse (no need to actually click on the square), a small grey dot will appear in the center and that is the center of the droplet that is displayed in that particular square.
48. Manually, drag each square to either the positive or the negative bin, depending on what you define as positive and negative.
49. If in doubt whether or not the droplet is positive or negative, you can either:
    a. Chose to delete it by left click of the mouse on the square and then "remove selected"
    b. Or, go over the square with the mouse and see where the center of the droplet because this should help you to see the actual region CPA has defined as the droplet.
    c. If you have another class, you want to sort it out as well (e.g. another fluorescence dye). Thus, you can add another classification bin through "Ad new class".
50. Once finished with the sorting, click on "Fetch!" again to get a new batch of droplets to sort.
51. When you are satisfied with your classification and you feel like you have sorted enough droplets for Classifier to distinguish the two groups correctly, click on "**Train**".
52. Based on your sorting, Classifier will automatically create 5 rules/features for classifying droplets.
53. You can add more rules/features in "Display", but, 5 should be enough for now. Also be aware of that SQLite databases have a limitation for how many rules/features that can be implemented.
    a. If you have 2 classes (as we have here – Positive/Negative) no more than 23 rules can be used
    b. For 3 classes, no more than 18 rules can be used etc.
54. Then click on "**Evaluate**". This creates a **Classification Report** that calculates cross-validation metrics based on the sorting you have done. Values closer to 1 indicate better performance of your sorting.
55. To evaluate based on a **Confusion Matrix**, instead choose *Evaluation > Confusion Matrix* and then click "**Evaluate**" again. This will create a matrix, where accuracy of your model is tested. Basically, it tells you how accurate the model classifies droplets based on the training set you have provided. Percent value closer to 100% indicate better accuracy.
56. Based on the evaluation, you can choose to (a) sort more droplets or (b) let Classifier try and score an image in your pipeline based on the model you have, to see if you have trained Classifier correctly by clicking on "Score Image" and then choosing an image number.
57. Two new windows will appear: (a) table with the data and (b) the image with illustration of which droplets were classified as positive (blue dot/mini-square) and negative (orange dot/mini-square).
58. If you are satisfied with the result, then you can:
    a. "Score All" images
    b. If you analyze many images at once, then you check the box where it says "*report enrichments?*" since this gives you more details regarding the scoring.
    c. Save your training set for later analysis in *File > Save Training Set.*
    d. Save your classification model for later analysis in *File > Save Classifier Model*.
